# ITPK1 regulates jasmonate-controlled root development in *Arabidopsis thaliana*

**DOI:** 10.1101/2023.06.16.545325

**Authors:** Naga Jyothi Pullagurla, Supritam Shome, Ranjana Yadav, Debabrata Laha

## Abstract

Jasmonic acid (JA) is a plant hormone that regulates a plethora of physiological processes including immunity and development and is perceived by the F-Box protein, Coronatine-insensitive protein 1(COI1). The discovery of inositol phosphates (InsPs) in the COI1 receptor complex highlights their role in JA perception. InsPs are phosphate-rich signaling molecules that control many aspects of plant physiology. Inositol pyrophosphates (PP-InsPs) are diphosphate containing InsP species of which InsP_7_ and InsP_8_ are the best characterized ones. Different InsP and PP-InsP species are linked with JA-related plant immunity. However, role of PP-InsP species in regulating JA-dependent developmental processes are poorly understood. Recent identification of ITPK1 kinase responsible for the production of 5-InsP_7_ from InsP_6_ *in planta* provides a platform to interrogate possible involvement of ITPK-derived InsP species in JA-related plant development. Herein this study, we report that ITPK1-defective plants exhibit increased root growth inhibition to bioactive JA treatment. The *itpk1* plants also show increased lateral root density when treated with JA. Notably, JA treatment does not induce ITPK1 protein level. Gene expression analyses revealed that JA-biosynthetic genes are not differentially expressed in the ITPK1-deficient plants. We further demonstrate that genes encoding different JAZ repressor proteins are severely downregulated in the ITPK1-defective plants. Taken together, our study highlights the role of ITPK1 in regulating JA-dependent root architecture development through controlling expression of different JAZ repressor proteins.

## 1. Introduction

Inositol phosphates (InsPs) are phosphate containing cellular messengers that control a large array of physiological processes in eukaryotes [1-4]. Combinatorial action of different kinases and phosphatases results into diverse inositol phosphate messengers of which InsP_6_ is one of the most abundant InsP species. InsP_6_, also known as phytic acid, contributes to different cellular processes either directly or indirectly by serving as a precursor for a class of signalling molecules known as inositol pyrophosphates (PP-InsP). PP-InsPs comprise of one or more diphosphate groups and are critical second messengers in yeast, amoeba and metazoan [5-7].

PP-InsP biosynthetic pathway is well established in yeast and mammals, where Kcs1/IP6K-type proteins synthesize 5-InsP_7_, and Vip1/PPIP5K-type kinases phosphorylate InsP_6_ and 5-InsP_7_ generating 1-InsP_7_ and 1,5-InsP_8_, respectively [8-12]. While Kcs1/IP6K-type proteins are absent in plants, Vip1/PPIP5K-type kinases are encoded by all plant genomes [13-15]. Similar to yeast and mammals, the *Arabidopsis* PPIP5K isoforms VIH1 and VIH2 catalyze the synthesis of InsP_8_ and are likely involved in synthesizing 1/3-InsP_7_ *in planta* (Figure 1) [14-17]. Recently, ITPK1 and ITPK2 were identified to catalyze the synthesis of 5-InsP_7_ from InsP_6_ *in vitro* [18-20] and *in planta* [17, 21, 22]. Identification of the proteins controlling PP-InsP synthesis *in planta* created new avenues to understand the physiological processes regulated by PP-InsP. These energy-rich species are emerging as signalling molecules in plants that are involved in regulating a plethora of cellular processes including nutrient sensing and immunity in plants [14-16, 23-30].

**Figure 1.**
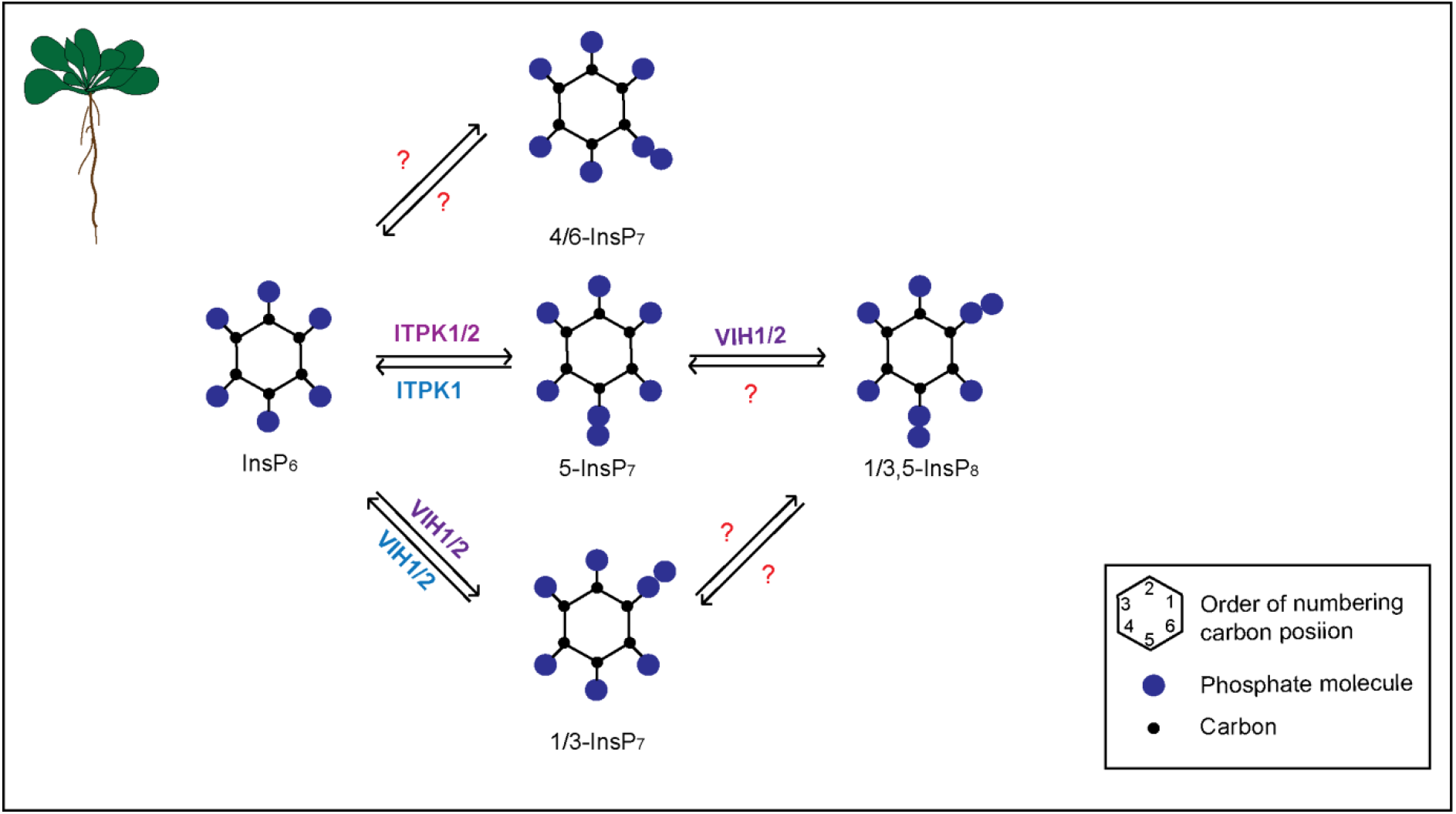
Inositol pyrophosphate (PP-InsP) synthesis in *Arabidopsis thaliana*. ITPK1/2 phosphorylates InsP_6_ at position C5 to generate 5-InsP_7_. VIH1/2 phosphorylates InsP_6_ and 5-InsP_7_ leading to the production of 1/3-InsP_7_ and 1/3,5-InsP_8_. Both ITPK1 and VIH1/2 can also catalyse the reverse reaction. Protein(s) responsible for the synthesis of 4/6-InsP_7_ in plants are not known currently and hence denoted by question mark.

Multiple studies have provided evidence that PP-InsPs play a regulatory role in the hormone signaling pathways including auxin, salicylic acid and jasmonic acid in *Arabidopsis* [14, 25, 28, 22]. Jasmonic acid (JA) is a phytohormone involved in plant development and immunity [31]. Jasmonoyl-isoleucine (JA-Ile), the bioactive JA-derivative, is perceived by the Coronatine Insensitive 1 (COI1) receptor protein that plays a crucial role in regulating JA responses [32-35]. Binding of JA-Ile to COI1 facilitates its interaction with different Jasmonate ZIM Domain (JAZ) transcriptional repressor proteins and promotes degradation, which derepresses MYC2 and other transcription factors, facilitating JA-dependent gene expression [36-41]. The crystal structure of the ASK1-COI1 receptor complex revealed distinct electron densities in the core of the solenoid structure [42]. These electron densities were likely due to individual phosphates that replaced an inositol phosphate ligand derived from the insect cells, possibly due to the high concentrations of ammonium phosphate used during crystallization [42]. Indeed, the nano-electrospray mass spectroscopy of insect-purified ASK1-COI1 protein confirmed the presence of an InsP_5_ species in the receptor complex [42]. Previous studies have implicated the role of inositol phosphates in plant wound response and disease resistance [43, 44]. Heterologous expression of human inositol 5-phosphatase in *Arabidopsis* has resulted in altered defense gene expression and increased weight of herbivorous caterpillar [45, 46]. The InsP_6_ defective mutant plants, *ipk1-1* and *ips2* exhibit susceptibility to various pathogens including the necrotrophic fungi *Botrytis cinerea* [44]. Moreover, investigating the *Arabidopsis ipk1-1* mutant revealed its increased sensitivity to JA [47]. Additionally, the *ipk1-1* mutant exhibited enhanced defense capabilities against herbivorous insects compared to the wild-type plants [47]. Collectively, these studies provide substantial evidence supporting the involvement of inositol phosphates in the regulation of JA-dependent responses in plants. Notably, *ipk1-1* plants also exhibit increased level of InsP_5_ [2-OH] and severely reduced levels of InsP_7_ and InsP_8_ species [14, 26, 27, 17].

To comprehensively assess the potential role of PP-InsPs on JA-dependent responses, consequences of VIH2 loss-of-function with impaired InsP_8_ synthesis in plants were monitored [14]. Although the *vih2* mutant plants exhibited unaltered levels of InsP_5_ [2-OH], they were susceptible to different insect herbivores and fungal necrotrophs [14, 25]. Furthermore, *vih2* mutants exhibited reduced JA-dependent gene expression despite elevated JA levels. Thus, the compromised resilience of *Arabidopsis vih2* mutants against herbivorous insects could be attributed to impaired JA perception rather than compromised JA production [14]. Furthermore, *in vitro* binding experiments using various radiolabelled InsPs revealed that higher inositol polyphosphates, namely InsP_6_ and InsP_7_, exhibit greater efficacy in binding to the ASK1-COI1-JAZ1-JA-receptor complex compared to lower InsPs [14, 25]. In conclusion, it has been proposed that coincidence detection of both VIH2-dependent InsP_8_ and jasmonate, form ASK1-COI1-JAZ receptor complexes that activate the JA-dependent gene expression [14]. Future works awaits to explore possible function of other InsP and PP-InsP species in JA-dependent physiological processes.

Analyses of *itpk1-2* mutant plants with altered level of various InsP and PP-InsP species [26, 17, 22] highlight a critical role of ITPK1 in different auxin-related processes including leaf venation, thermomorphogenic responses, and primary root elongation [22]. Notably, JA also controls root growth and development [48-50]. Since ITPK1 is responsible for the production of 5-InsP_7_, a precursor of InsP_8_ and that InsP_8_ is linked with JA-perception, further investigation is necessary to elucidate the potential involvement of ITPK1-derived inositol phosphate species in JA-mediated physiological processes. Herein this study, we show that ITPK1 activity is critical for jasmonate-dependent root development.

### 2. Materials and Methods

#### 2.1. Arabidopsis Plant Material and Growth Conditions

Seeds of T-DNA insertion lines of *Arabidopsis thaliana* (ecotype Col-0) were obtained from the Arabidopsis Biological Resource Center at Ohio State University (http://abrc.osu.edu). The *itpk1-2* plants and the *itpk1-2* transgenic lines expressing the genomic *ITPK1* fragment used in this study were reported previously [22]. Wild-type and all relevant transgenic lines were amplified together on soil and perlite mix under identical conditions (16-h light and 8-h dark, day/night temperatures 22/18°C and 120 µmol^-1^ m^-2^ light intensity). For sterile growth, seeds were surface sterilized in solution containing 70% (v/v) ethanol and 0.05% (v/v) Triton X-100 for 10 min and washed twice with 90% (v/v) ethanol. Sterilized seeds were sown on solidified half-strength Murashige and Skoog (MS) media (0.8% phytagel), supplemented with 1% sucrose, and then stratified for 2 days at 4°C, and grown in a Percival plant chamber under conditions of 8-h light (22°C) and 16-h dark (20°C).

#### 2.2. Primary Root Length and Lateral Root Density Assay

For primary root length and lateral root density measurement, seedlings were grown vertically on solidified half-strength MS media, supplemented with 1% (w/v) sucrose under 8-h light/16-h dark condition. After 7 days of growth, the seedlings were transferred on to modified half-strength MS media supplemented with 1% (w/v) sucrose and with or without 50 µM MeJA. The primary root length was noted at intervals of 3, 5 and 7 days. Images were taken using Bio-Rad ChemiDoc. The primary root length and lateral roots were quantified using ImageJ software.

#### 2.3. RT-PCR Analyses

9-day-old seedlings were harvested for total RNA extraction using TRIzol reagent (Sigma). Then, 1 µg of total RNA was used to synthesize cDNA with PhiScript™ cDNA Synthesis Kit (dx/dt). The cDNA samples were diluted to 150 ng μL^−1^ of which 0.5 uL was used as template for reaction. The qPCR was performed using the DyNAmo ColorFlash SYBR Green qPCR Kit (Thermo-scientific) with CFX96 Touch Real-Time PCR Detection System (Bio-Rad) according to the manufacturer’s protocol (Bio-Rad). The relative quantitation method (^ΔΔ^CT) was used to evaluate quantitative variation among replicates. *Beta-TUBULIN* was used as reference gene.

#### 2.4. Western analyses

For immunoblot analyses, 9-day-old seedlings of *itpk1-2* lines expressing genomic ITPK1 fragment with C-terminal G3GFP fusion (compl. line#7) were grown on sterile solidified [0.8% (w/v) phytagel], half-strength MS media, supplemented with 1% (w/v) sucrose for 7 days, then transferred to liquid half-strength MS media (pH 5.7), supplemented with 1% (w/v) sucrose and with or without 50 µM MeJA, and subsequently incubated for indicated time points before harvesting. Seedlings were homogenized (w/v) in protein extraction buffer [5 mM Tris–HCl, pH 7.5, 150 mM NaCl, 10 mM MgCl_2_, 1 mM EDTA] containing 1x plant protease inhibitor (Sigma). The extract was centrifuged at high spin for 10 min at 4°C. The supernatant was used for western blot analysis. Protein extracts were boiled with loading buffer containing SDS for 10 min, and proteins were separated by SDS-PAGE and detected with anti-GFP antibodies (Roche, product No. 11814460001).

## 3. Results

### 3.1. Arabidopsis itpk1-2 Lines Exhibit Enhanced Jasmonate-Mediated Root Growth Inhibition and Increased Lateral Root Formation

To investigate whether JA-mediated processes are altered in ITPK1-deficient plants, we monitored root development [40-48] and lateral root (LR) formation [50] after exogenous application of methyl jasmonate (MeJA), a bioactive form of jasmonate [49]. The impact of MeJA treatment on root development was assessed for wild-type, *itpk1-2* and *itpk1-2*::*AtITPK1G3GFP* complementary lines (Figure 2). Notably, primary root growth inhibition was more pronounced in *itpk1-2* plants when compared with wild-type seedlings (Figure 2a, b, c and d). Furthermore, LR density (Figure 2e) was significantly increased in *itpk1-2* plants than wild-type plants after MeJA treatment. Importantly, the altered root architecture of ITPK1-defective plants was rescued in the *itpk1-2* lines expressing the ITPK1-G3GFP translational fusion under the control of its endogenous promoter. Collectively, these data suggest that *itpk1-2* plants are hypersensitive to JA and are defective in JA-dependent root architecture development.

**Figure 2.**
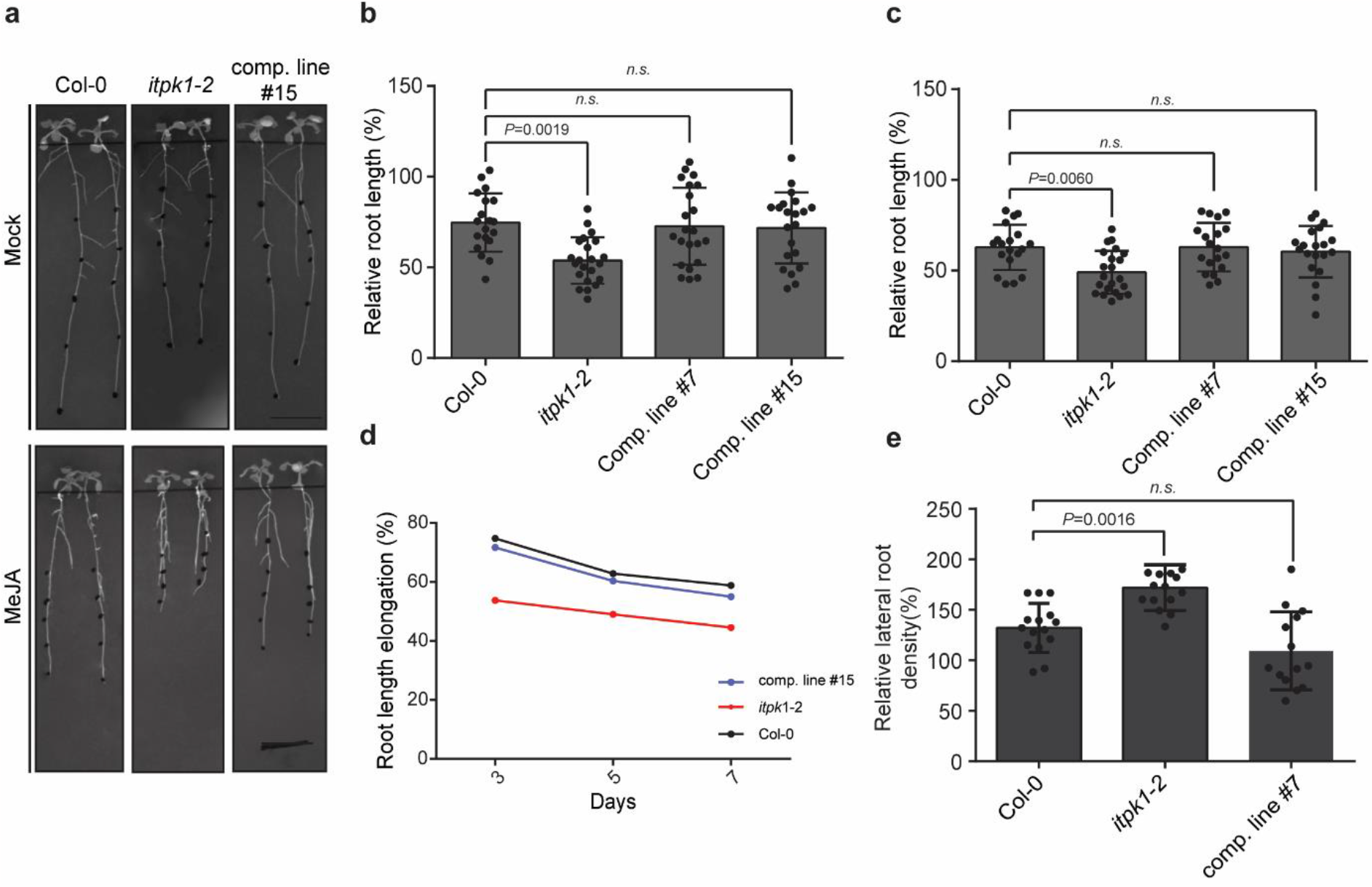
Sensitivity of *itpk1-2* plants to methyl jasmonate (MeJA) treatment. (**a**) Representative pictures of seedlings of wild-type (Col-0), *itpk1-2* and one complemented *itpk1-2* line grown in presence of mock and MeJA. Seeds were germinated on solidified half-strength MS agar media, supplemented with 1% (w/v) sucrose. After 7 days, seedlings were transferred to solidified half-strength MS media supplemented with 1% (w/v) sucrose and 50 µM MeJA. The percentage of changes in primary root length induced by 50 µM MeJA was determined for the wild-type, *itpk1-2* mutant, and the selected complemented lines. The growth of root length was evaluated after (**b**) 3 days and (**c**) 5 days using ImageJ software. The data presented are means ± SD (*n* ≥ 20). The *P* values depict the significance in two-way ANOVA followed by Tukey’s test. (**d**) The relative root elongation of the designated genotypes under 50 µM MeJA treatment. Different letters indicate significance in two-way ANOVA followed by Tukey’s test (d and a, *P* < 0.01; e and b, *P* < 0.05; f and c, *P* < 0.05). The experiment was repeated multiple times with consistent results. (**e**) The lateral root density of the designated genotypes was quantified after 50 µM MeJA treatment. Lateral roots were quantified using ImageJ software. Images were taken using Bio-Rad ChemiDoc. The data presented are means ± SD (*n* ≥ 20). The *P* values depict the significance in two-way ANOVA followed by Tukey’s test.

#### 3.2. Effect of Methyl Jasmonate (MeJA) on ITPK1 Protein Level and in the Expression of Several Jasmonate-Biosynthetic Genes

Since *itpk1-2* plants show altered JA response, we wanted to investigate whether MeJA treatment could influence ITPK1 expression. Our findings revealed that there was no observable difference in the content of the ITPK1 protein after MeJA treatment (Figure 3a and b). These results indicate that MeJA does not exert regulatory control over the expression of ITPK1 in plants.

**Figure 3.**
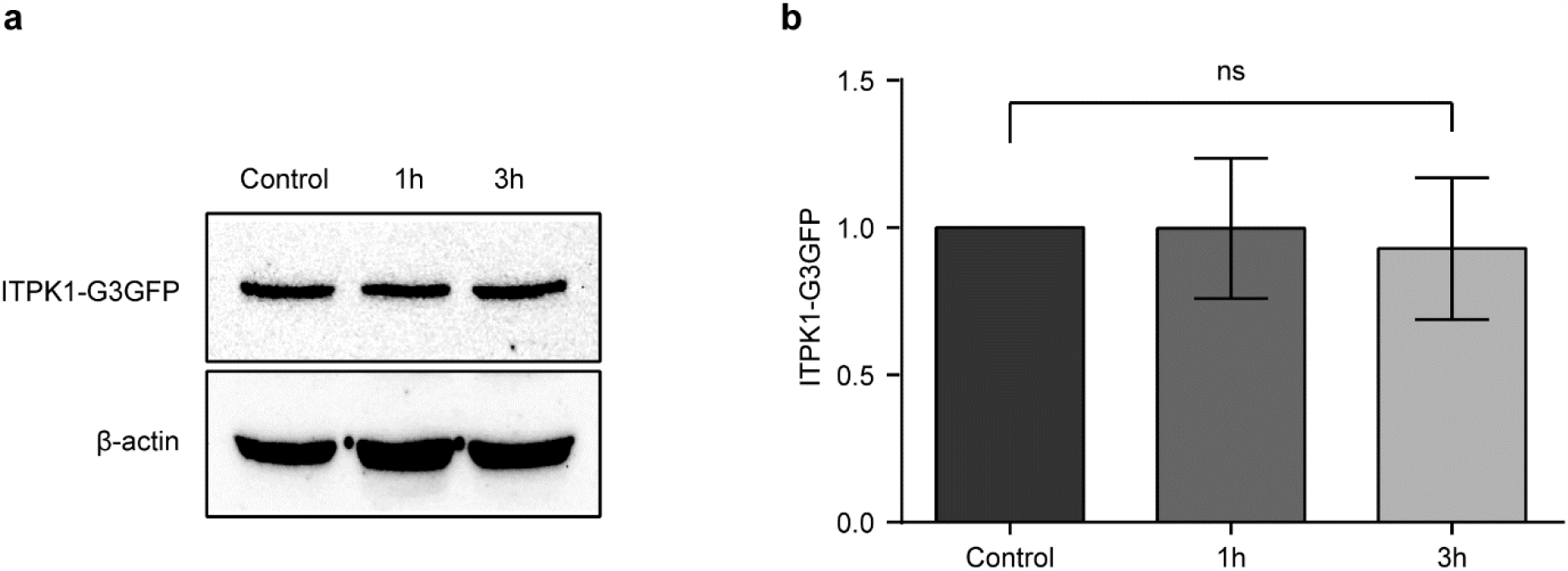
Effect of methyl jasmonate (MeJA) on the expression of ITPK1 protein. (**a**) Western blot analysis of total protein extract prepared from 9-day-old compl. line#7 seedlings expressing ITPK1 in translational fusion with N-terminal G3GFP. The 9-day-old seedlings were transferred to liquid half-strength MS media (pH 5.7), supplemented with 1% (w/v) sucrose and with or without 50 µM MeJA, and then incubated for indicated time points before harvesting. Approximately 25-30 µg of protein was loaded in each well, and ITPK1 was detected with antibodies against GFP (Roche). The β-actin (45 kDa) was used as an internal loading control. (**b**) Quantification of ITPK1 protein levels. Graph showing the relative ITPK1 protein levels following treatment of MeJA for 1-h and 3-h with β-actin as control. Relative ITPK1 protein levels were quantified using ImageJ software. The data presented are means ± SD (*n* = 3). The ns depict the non-significance in two-way ANOVA followed by Tukey’s test. The experiments were repeated three times with similar results.

Further to get mechanistic insight on role of ITPK1 in JA-mediated root development, we conducted qPCR analyses to study the expression levels of various genes related to biosynthesis and perception, namely *ALLENE OXIDE SYNTHASE* (*AOS*), *JASMONATE RESISTANT 1* (*JAR1)*, and *COI1* in wild-type and *itpk1-2* plants. These lines were subjected to treatment with 50 µM MeJA. No significant differences were observed in the expression level of any of these genes between the wild-type and *itpk1-2* plants (Figure 4). Collectively, these results indicate that ITPK1 does not play a role in modulating the expression of these genes during JA-dependent root development.

**Figure 4.**
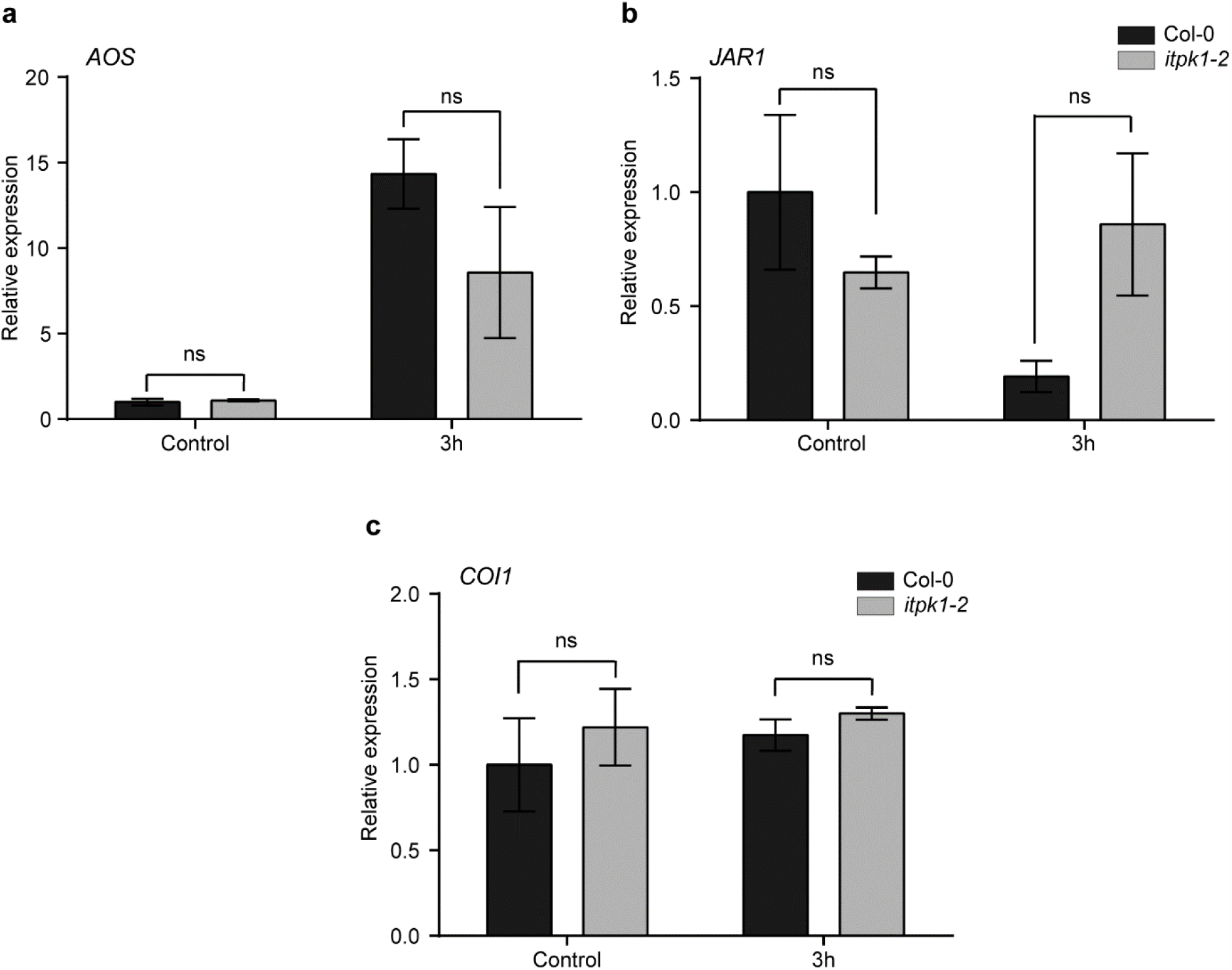
The relative expression levels of jasmonate-related genes (**a**) *AOS* (**b**) *JAR1* and (**c**) *COI1* were assessed in seedlings of the wild-type and *itpk1-2* mutant. 7-day-old seedlings were harvested at specified time points following methyl jasmonate (MeJA) application, along with the untreated plants. The reference gene used for normalization was *β-TUBULIN*. The results are presented as means ± SD (*n* = 3). The significant differences between the lines for each gene, determined through two-way ANOVA and Tukey’s test.

### 3.3 ITPK1 Activity is Required for the Regulation of Methyl Jasmonate (MeJA) Induced JAZ Transcript Levels

JAZ proteins are critical repressors of jasmonate signaling [36, 37, 42]. Further, the *jazQ* mutant seedlings lacking five JAZ repressors show an increased root growth inhibition to exogenous MeJA treatment when compared with wild-type seedlings [51]. Hence, we delved into understanding whether the absence of ITPK1 would affect the expression of genes encoding different JAZ *repressors*. As anticipated, the wild-type plants exhibited increased *JAZ* expression after MeJA treatment, whereas *itpk1-2* plants showed a robust decrease in expression of various *JAZ* genes, namely *JAZ1, JAZ2, JAZ5* and *JAZ9* (Figure 5). These findings suggest that ITPK1 activity is critical for MeJA-dependent *JAZ* expression.

**Figure 5.**
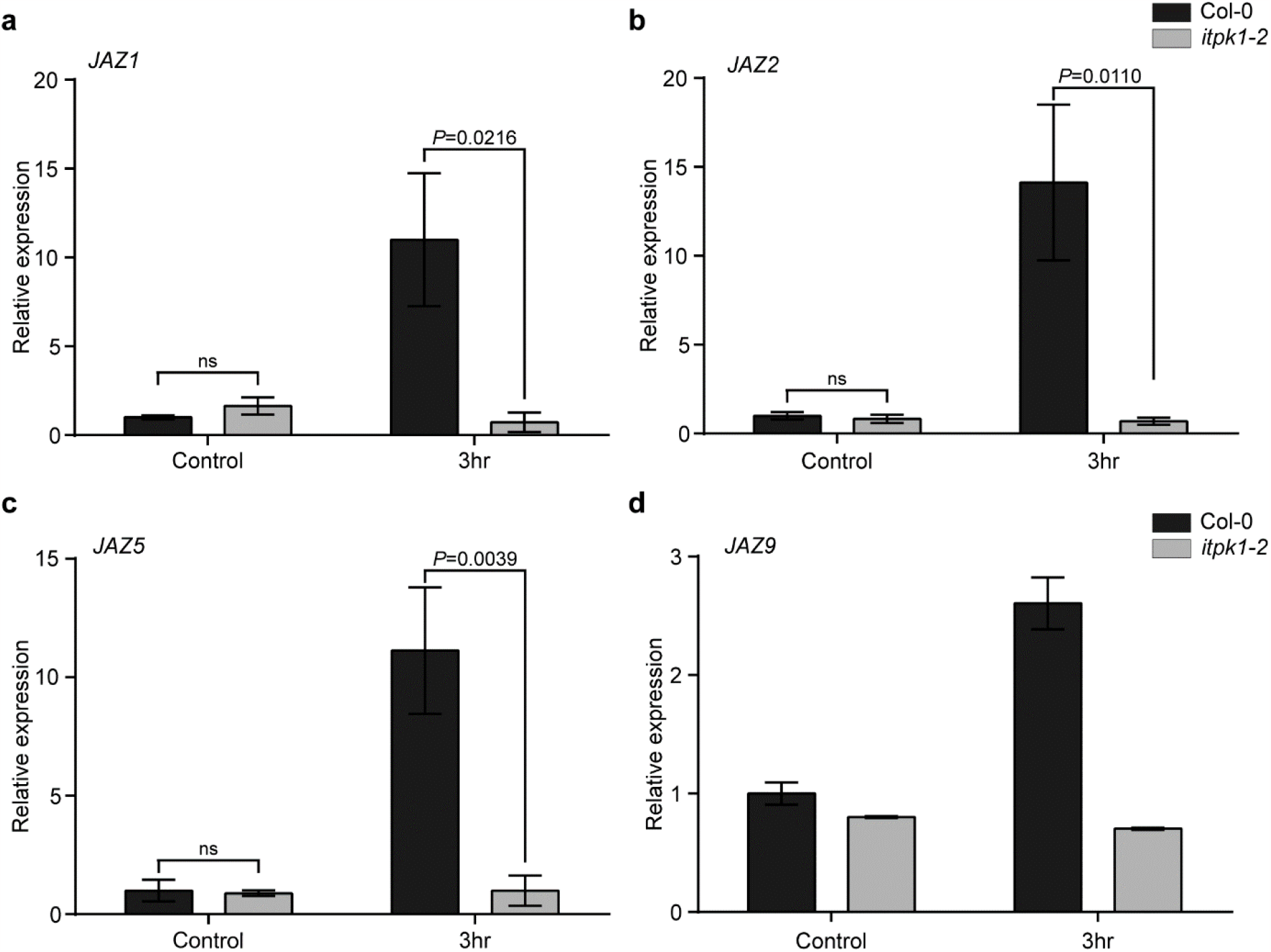
ITPK1 controls MeJA-dependent expression of (**a**) *JAZ1* (**b**) *JAZ2* (**c**) *JAZ5* and (**d**) *JAZ9* genes. 7-day-old seedlings were harvested at specified time points following methyl jasmonate (MeJA) application, along with the untreated plants. The relative transcript abundance was calculated by normalizing to the reference gene *β-TUBULIN*. The data presented as means ± SD (*n* = 3). Data were analyzed by a two-way ANOVA using Tukey’s test. The qPCR analyses were repeated with similar results.

## 4. Discussion

Jasmonic acid is a critical signaling compound that regulates varied physiological processes, such as root growth inhibition, anthocyanin accumulation, and stress responses [31, 52-55]. Although the inositol phosphates have been shown to regulate jasmonate signaling pathways, it is still largely unclear whether a specific InsP isomer mediates certain JA responses or different InsP species work collaboratively to modulate specific JA processes in plants.

In this current study, we demonstrate that function of ITPK1 is critical for jasmonate signaling. Considering the enhanced lateral root formation and sensitivity of *itpk1-2* mutant roots to MeJA, we propose that ITPK1 acts as a negative regulator in the JA-dependent root development. Our experiments suggest that ITPK1 level remains unaltered after MeJA treatment however further work is required to unravel any post-translational modification of ITPK1 after the hormone treatment. Jasmonate signaling entails extreme transcriptional reprogramming involving intricate interactions between both positive and negative regulators. The JAZ repressors are responsible for modulating the jasmonate signaling by interacting with various transcription factors [52, 56-59]. Compromised expression of various *JAZs* after MeJA treatment in the ITPK1-deficient plants highlights the importance of ITPK1 activity in maintaining cellular *JAZ* levels. Overall, these findings further suggest that hypersensitive phenotype to MeJA of *itpk1-2* mutant could be due to reduced *JAZ* expression in these mutant plants. In conclusion, the findings of this study provide insights into the role of ITPK1 in JA-mediated root development. Further studies are required to understand the mechanism underlying the hypersensitive phenotype of *itpk1-2* on MeJA treatment. It will be interesting to examine the possibility of different inositol phosphates forming a series of distinctive JA co-receptor complexes and the physiological importance of such diverse co-receptor complexes in plants.

## Author Contributions

D.L. designed and supervised the research. N.J.P and S.S. performed all the experiments. N.J.P., S.S., R.Y. and D.L. analyzed the data. N.J.P. and D.L. wrote the manuscript with input from all the authors.

## Funding

This work was supported by grants from the Science and Engineering Research Board (SERB) SRG/2021/000951, the Department of Biotechnology (DBT) for HGK-Innovative Young Biotechnologist Award (BT/13/IYBA/ 2020/04), and Indian Institute of Science start-up fund to D.L. N.J.P. acknowledges Council of Scientific & Industrial Research (CSIR) for research fellowship. S.S. and R.Y. are recipient of IISc research fellowship.

## Institutional Review Board Statement

Not applicable.

## Informed Consent Statement

Not applicable.

## Data Availability Statement

The data that support the findings of this study are all provided within the body of the manuscript.

## Notes

### Competing Interest Statement

The authors have declared no competing interest.

